# Noise-induced changes in rat head stability and orientation correlate with dysfunction of otolith irregular afferents

**DOI:** 10.1101/2023.10.26.563800

**Authors:** Mamiko Niwa, David Bauer, Marie Anderson, Ariane Kanicki, Katharine Keller, Richard A. Altschuler, Courtney E. Stewart, W. Michael King

**Affiliations:** Kresge Hearing Research Institute, University of Michigan; VA Ann Arbor Healthcare System

## Abstract

Exposure to intense noise damages both the cochlea and vestibular end organs. Our group previously reported attenuated vestibular short-latency evoked potentials (VsEP) and reduced numbers of calretinin-positive (CR+) calyces in the saccule following noise exposure. Here, we examined rats’ resting head orientation with respect to gravity as well as head stability following a 4-hour exposure to 120 dB SPL noise. We also assessed how behavioral changes are related to changes in VsEP waveforms and calretinin expression in the utricle and saccule to elucidate potential underlying mechanisms. We found significant reductions in the P2N2 and N2P3 amplitudes following noise exposure. The number of CR+ calyces in both saccule and utricle were also significantly reduced. The size of the reduction in N2P3 amplitude was significantly correlated to the number of CR+ calyces. Animals with a greater loss of CR+ calyces in the utricle following noise showed significant decreases in the average speed of y-axis rotational head motion, while those with a fewer loss of CR+ calyces showed significant increases. In addition, animals with larger noise-induced changes in VsEP and CR+ calyces held their heads motionless longer following noise exposure. We hypothesize that noise exposure is inherently destructive to an animal’s head stability and thereby manifests as an increase in average head speed in mildly to moderately affected animals. But when the damage was large enough, animals exhibited reduced duration and head motion speed as a behavioral adaptation. The noise exposure also significantly altered the pitch angle of head orientation in animals who had the largest reduction in CR+ calyces in the saccule, suggesting that saccular irregular afferents, including those that are CR+, are critical in control of head and body posture.

## Introduction

Exposure to intense noise has been widely accepted as a hazard to our hearing, but less accepted is the effect of noise exposure on the well-being of vestibular function. Impairment of the vestibular system, whose canonical functions include stabilization of gaze, head motion, and posture, can lead to debilitating symptoms such as vertigo, blurred vision during head movement and postural instability. Furthermore, inputs originating from the vestibular periphery diffusely innervate various cortical areas and the limbic system; thus, vestibular dysfunction can have far-reaching impacts on various cognitive functions such as spatial navigation, memory, and attention (reviews in Shinder and Taube, 2010; Smith and Zheng, 2013). Because vestibular impairments are especially prevalent among the elderly, it is important to identify external factors and underlying mechanisms that lead to dysfunction of the vestibular periphery.

Clinical studies of industry and military workers with noise-induced hearing loss (NIHL) has long noted concomitant abnormalities in the vestibular periphery, implicating noise exposure as damaging to both the auditory and vestibular organs in the inner ear (Collins, 1948). Abnormal nystagmus as well as reduced gain in the vestibulo-ocular reflex in the NIHL patients indicate dysfunction of the semi-circular canal organs, which encode angular acceleration of the head (Man et al., 1980; Oosterveld et al., 1982; Shupak et al., 1994). Hypofunction of the otolith organs, which encode linear acceleration of the head, was also indicated in the NIHL patients, as evidenced by reduced vestibular-evoked myogenic potentials of cervical and extraocular muscles (cVEMP and oVEMP; Akin et al., 2012; Tseng and Young, 2013; Viola et al., 2020; Wang and Young, 2007).

In animal models, noise exposure has been shown to induce molecular and morphological changes in the vestibular periphery including elevated oxidative stress markers (Tamura et al., 2012), reduced calcium-binding protein in afferent terminals (Stewart et al., 2018, 2020), hair bundle disruption (Stewart et al., 2016), vacuole formation and loss of nucleus in type-I hair cells (Hsu et al., 2008), degeneration of epithelial cells (Akdogan et al., 2009), and structural collapse and/or membrane rupture (Mangabeira-Albernaz et al., 1959; Mccabe and Lawrence, 1958) depending on the intensity, duration, and frequency profiles of the noise exposures (review by Stewart et al., 2020). Electrophysiological changes were also noted as reduced amplitude of the cVEMP (Hsu et al., 2008; Kumar et al., 2010) and reduced amplitude and prolonged latency in the vestibular short-latency evoked potential (VsEP; Perez et al., 2002; Stewart et al., 2018, 2020).

Behavioral or clinical consequence of noise-induced damage to the vestibular periphery is less well understood. Notably, among the NIHL patients with abnormal findings in the vestibular periphery, only a subset complained of symptoms such as vertigo and dizziness. However, the absence of clinical symptoms does not mean there is no behavioral consequence; studies have shown that the NIHL patients without clinically relevant vestibular symptoms show larger body sways compared to control subjects (Juntunen et al., 1987; Ylikoski et al., 1988). Animal studies also show that noise exposure reduced rodents’ proficiency in motor tasks that require balance (rota-rod: Tamura et al., 2012; balance beam crossing: Bartikofsky et al., 2023). Potential mechanisms underlying these behavioral outcomes include reduced gain and/or increased variability in encoding the gravitational vector, reduced accuracy in an internal model of active vs passive head movements, and reduced speed/gain in the vestibulo-collic and -spinal reflexes, degrading stability of head and body posture.

A goal of our study was to provide a quantitative analysis of behavioral changes that follow noise exposure. Here, we examined animals’ head orientation and stability during self-generated head movements before and after exposure to 120 dB noise. Our group recently reported that rats exposed to a similar noise profile exhibited significantly decreased proficiency in balance-beam crossing (Bartikofsky et al., 2023). We also showed that this noise exposure attenuated VsEP waveforms and reduced the number of countable calretinin-positive calyces in the saccule (Stewart et al., 2018, 2020). The present study identifies a relationship between electrophysiological and morphological changes in the periphery to changes in head stability and orientation and suggests potential underlying mechanisms of noise-induced vestibular dysfunction.

## Results

The vestibular short-latency evoked potential (VsEP) was measured before and after exposing rats to 3-octave band (0.53 to 4.2 kHz) noise at 120 dB SPL for 4 hours. The VsEP is thought to be a far field potential, generated by synchronous discharges of the action potentials of irregular vestibular afferents and vestibular neurons in the brain stem, and evoked by a linear acceleration jerk applied to the animal’s head (Nazareth and Jones, 1998). The rate of change in linear acceleration, jerk, has been shown as a parameter on which the VsEP primarily depends (Jones et al., 2011).

A representative animal’s VsEP responses to 5 jerk levels are shown in Figure 1A prior to noise exposure (black traces), 1 week (magenta traces) and 4 weeks after noise exposure (green traces). The VsEP typically starts with a positive peak, P1 (denoted with a grey arrow in Figure 1A), followed by multiple sets of negative and positive peaks (Jones and Jones, 1999; Plotnik et al., 1997). Additionally, we defined the N0 (a grey star) as the point of deflection leading to the P1 peak. In this animal, P1N1 amplitude, defined as the potential difference from P1 to N1 peak, was reduced at 1 and 4 weeks after noise compared to the baseline at all stimulus levels (2^nd^ panel from the left in Figure 1B). A reduction was also seen for the P2N2 and N2P3 amplitudes, but not for the N0P1 (Figure 1B). The P1 peak was delayed, while P3 peak occurred earlier, both at 1 and 4 weeks after noise for most stimulus levels (Figure 1C).

**Figure 1.**
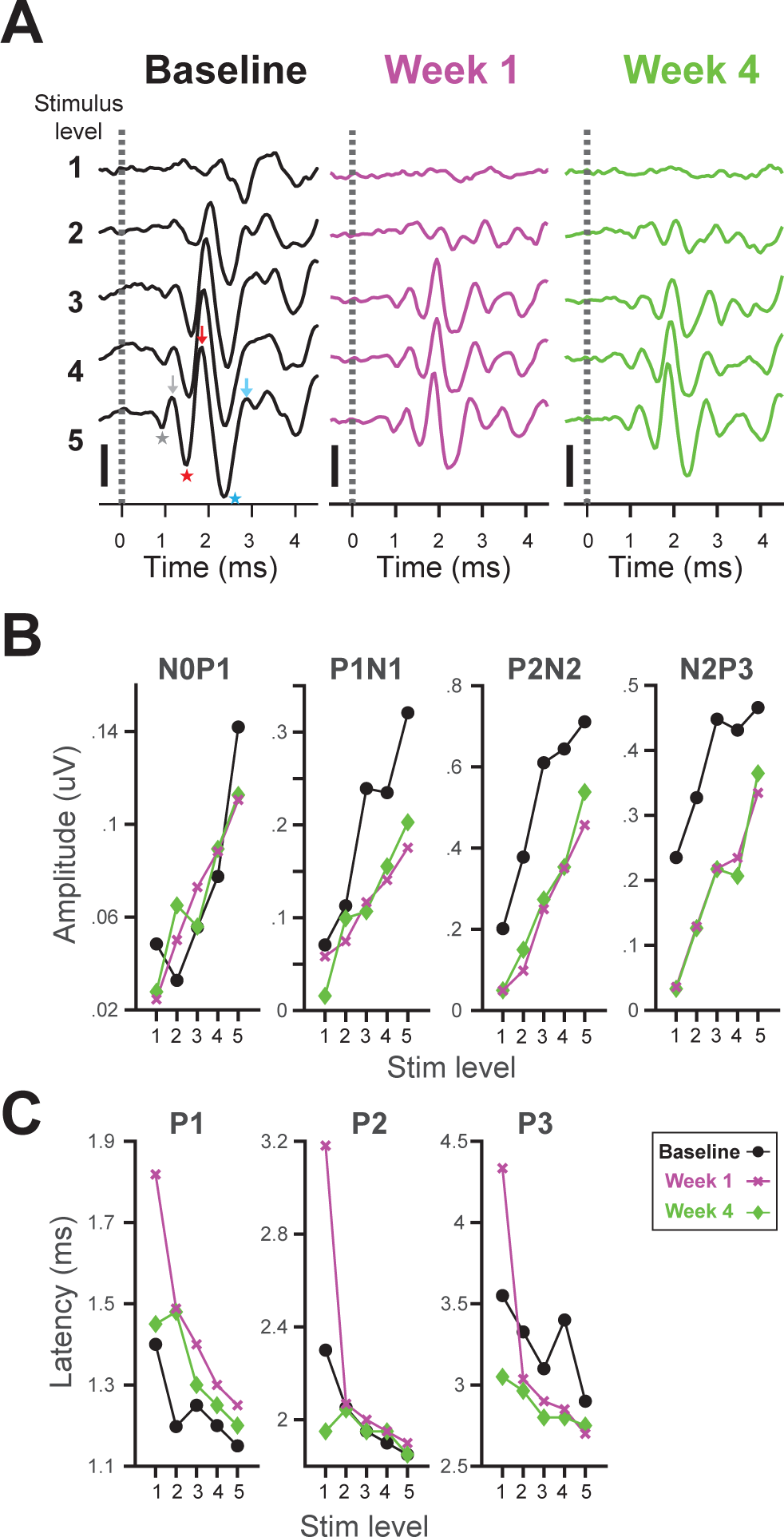
Noise exposure attenuates VsEP responses. **A**: Representative VsEP recordings before noise exposure (Baseline, black traces), 1-week post-exposure (magenta traces), and 4-week post-exposure (green traces). Stimulus jerk levels were (1) 0.32, (2) 1.1, (3) 2.2, (4) 3.2, and (5) 5.5 G/ms. VsEP peaks N0, P1, N1, P2, N2, and P3 peaks are denoted with a gray star, gray arrow, red star, red arrow, blue star, and blue arrow, respectively, on the left bottom VsEP trace. **B**: N0P1, P1N1, P2N2, and N2P3 amplitudes (from left to right) as a function of stimulus level at baseline (black), 1 week (magenta), and 4 weeks (green) after noise exposure. N0P1 amplitude is defined as the distance from N0 to P1 peak, and similarly for the other. **C**: P1, P2, and P3 peak latency as a function of stimulus level.

Across the population, the P2N2 and N2P3 amplitudes were significantly reduced at both 1 and 4 weeks after noise exposure compared to the baseline, while there was no significant change in the P1N1 amplitude due to large variability among animals (Figure 2A). The N0P1 amplitude was significantly increased at both 1 (p = 0.0020) and 4 weeks (p = 0.0003) after noise exposure when the change was collapsed across all stimulus levels (left most panel on Figure 2A). We found no significant change in the latencies of N0, N1, N2, P1, and P2 peaks, however, the P3 peak latency was significantly faster at all-levels-collapsed at week 4 (p = 0.0015; right panel in Figure 2B). This appears to be because the amplitude of the P3 component is smaller and peaking earlier after noise exposure.

**Figure 2.**
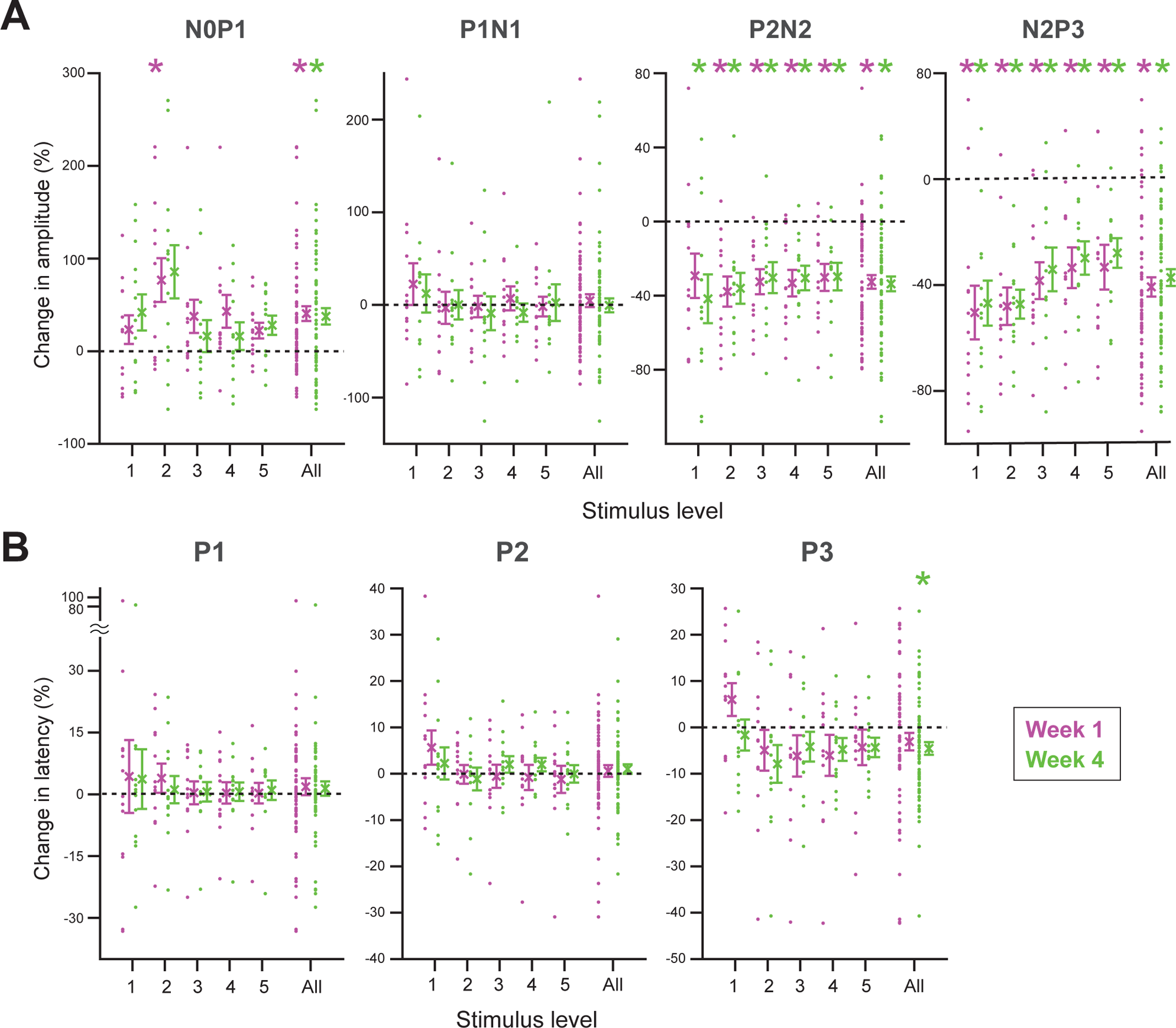
**A**: The % change in the N0P1, P1N1, P2N2, and N2P3 amplitudes (from left to right) at 1 week (magenta) and 4 weeks (green) after noise exposure compared to the baseline as a function of stimulus level. The % change values are collapsed across stimulus levels and shown at the column denoted as ‘All’ to show the overall effect. Each dot denotes an individual animal’s data point, the cross symbols denote population means, and error bars denote standard error of the mean (SEM). Asterisks denote significance levels at p < 0.01 and p < 0.05 by t-test for individual stimulus levels and all-levels collapsed, respectively. **B**: % change in P1, P2, and P3 peak latency as a function of stimulus levels.

### The reduced number of the calretinin-positive calyces in the utricle correlates with reductions in VsEP amplitude

Next, we examined the expression of a calcium-binding protein, calretinin, in the calyces of both the saccule and utricle at 5 weeks after noise exposure. Calretinin is expressed in the calyceal terminations of afferents contacting only type-1 hair cells (calyx-only afferents), but not in the terminations of afferents contacting only on type-2 hair cells (bouton afferents) or those contacting both type-1 and type-2 hair cells (dimorphic afferents; Desai et al., 2005; Desmadryl and Dechesne, 1992; Leonard and Kevetter, 2002). Afferents with calretinin-positive (CR+) calyces in the otoliths are known to have irregular temporal firing patterns and large gain and phase leads relative to head motion (Goldberg et al., 1990) and to be sensitive to high-frequency changes in linear acceleration of the head, such as the jerk stimuli that cause the VsEP.

We found that the number of CR+ calyces in the saccule and utricle of noise-exposed animals was significantly reduced compared to control animals (Figure 3A and 3B). The number of CR+ calyces in the utricle and saccule was significantly correlated with the degree of the N2P3 amplitude reduction at all but the smallest level at week 1 (stim level 5: r = 0.81, p = 0.00070, stim level 4: r = 0.83, p = 0.00047, stim level 3: r = 0.81, p = 0.00070, stim level 2: r = 0.68, p = 0.011; Figure 3C). The P3 peak latency at week 4 was significantly correlated with the number of CR+ calyces in the utricle, but not the saccule, at stim level 4 (r = 0.58, p = 0.036; Figure 3D). Thus, animals with a larger reduction in CR+ calyces is associated with a smaller P3 component peaking earlier compared to the baseline.

**Figure 3.**
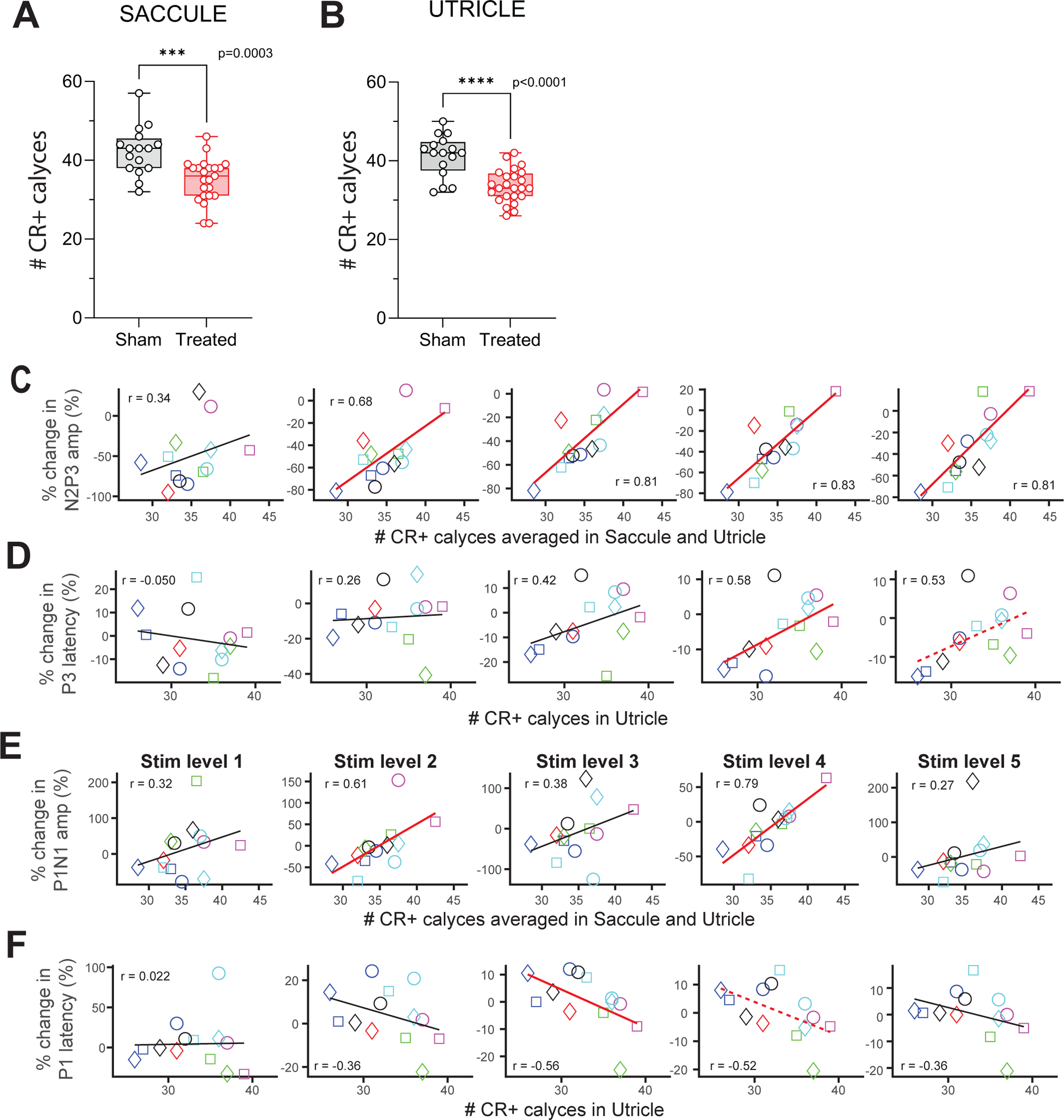
**A**: Number of CR+ calyces in the saccule of sham animals (black) and noise-treated animals (red). **B**: The same as **A**, but for the utricle. **C**: % change in N2P3 amplitude at 1 week after noise exposure for stimulus level 1 through 5 (from left to right panels) against the number of CR+ calyces averaged between the saccule and utricle. **D**: % change in P3 latency at week 4 for stimulus level 1 through 5 (from left to right panel) against the number of CR+ calyces in the utricle. **E**: Same as **C**, but for % change in P1N1 amplitude. F: Same as D, but for % change in P1 latency. In panels **C**∼**F**, linear fits are shown as a red solid line for significant correlations (p < 0.05) with Pearson’s correlation test. Dotted red lines denote marginally significant correlations (p < 0.10). Each combination of symbol and color designates an individual animal’s data point, and is consistently used for all other figures.

The P1N1 amplitude change was also significantly correlated with the number of CR+ calyces averaged between the utricle and saccule (stim level 4: r = 0.79, p = 0.0013, stim level 2: r = 0.61, p = 0.027; Figure 3E). The P1 peak latency was correlated the number of CR+ calyces in the utricle, but not the saccule (stim level 3: r = −0.56, p = 0.045; Figure 3F). Thus, animals with a larger reduction in CR+ calyces show a smaller and delayed P1N1 wave, while those with a smaller reduction in CR+ calyces show a larger and faster one compared to the baseline. Due to these opposing effects, the population-average change in the P1N1 amplitude and the P1 latency were diminished to 0 (Figure 2A, B).

### Animal’s head stability and orientation following noise exposure

Next, we present how these electrophysiological and morphological changes affected head stability and orientation following noise exposure. Using a miniature motion sensor attached to an animal’s head, we examined their head orientation and stability during inter-trial-intervals (ITIs) between abrupt, horizontal rotations given in our behavioral experiment (see Methods for behavioral test). During the ITIs, no external motion was applied to the rat, and it could freely move its head. The present study includes behavioral changes following exposure to sodium arsanilate, a vestibulo-toxic compound, as a positive control of bilateral vestibular loss (see Methods for trans-tympanic injection of sodium arsanilate).

### Animals with a greater noise-induced damage keep their head motionless longer following noise exposure

To assess head stability, we first quantified how long an animal’s head was stationary during ITIs. Figure 4A shows ‘stable period duration’, measuring how long each stationary interval lasted, over the course of the behavioral experiment in a representative noise-exposed animal. The daily median of stable period duration significantly increased after noise exposure (p = 3.4 x 10^−4^ with rank-sum test, Figure 4B). Figure 4D and 4E show that within each ITI, the total amount of time an animal’s head remained stationary, ‘the proportion of stable time’, also increased after noise exposure in this animal (p = 2.4 * 10^−4^, Figure 4E). We found that a subset of noise-exposed animals significantly increased stable period duration (5 out of 17 animals, Figure 4C) and proportion of stable time (2 out of 17 animals, Figure 4F), suggesting that noise exposure resulted in an apparent promotion of head motionlessness in these animals.

**Figure 4:**
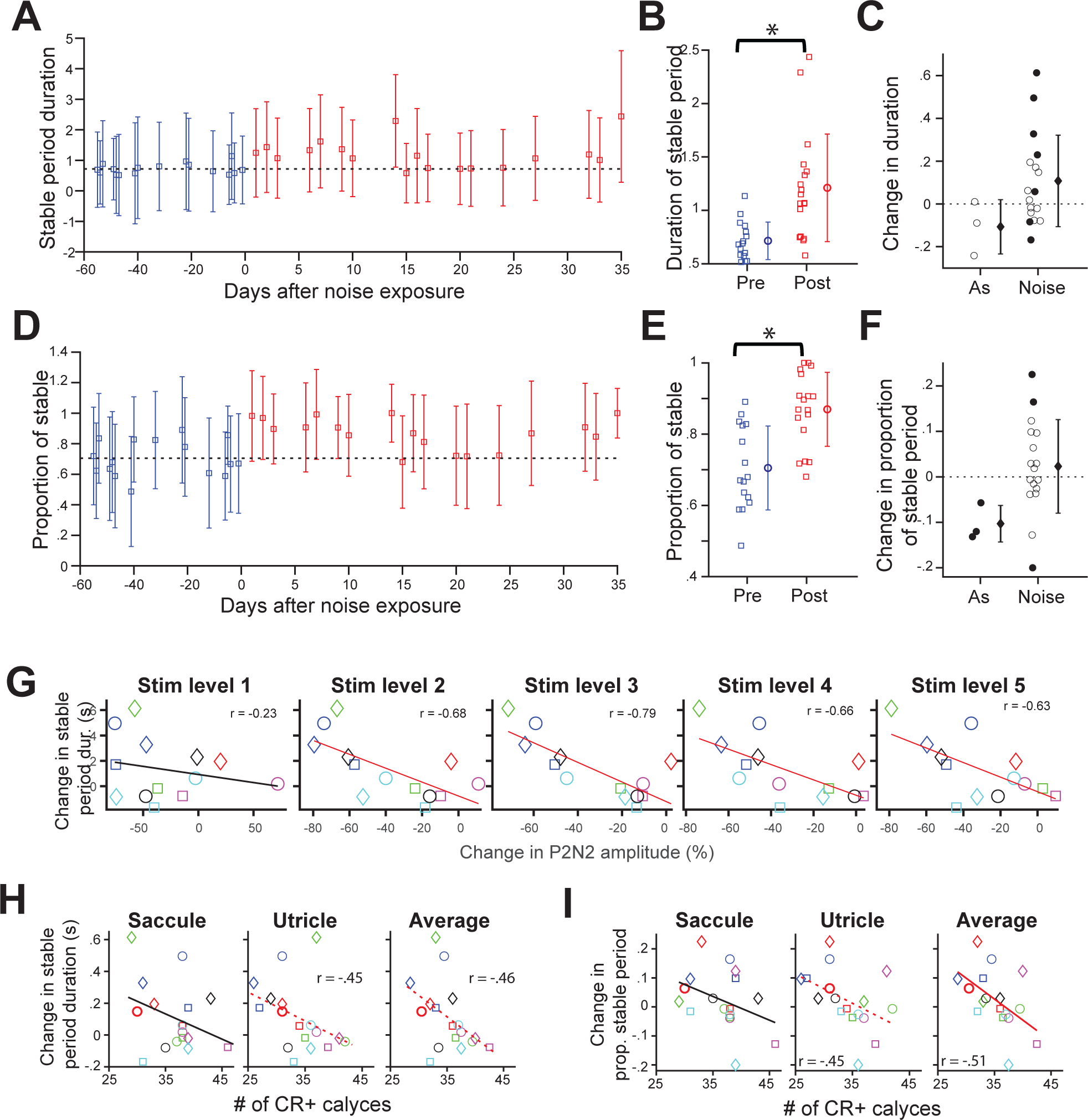
Noise exposure causes reduced head motion durations in relation to VsEP changes and numbers of reduced CR+ calyces in the otolith organs. **A**: The stable period duration versus days before or after noise exposure for a representative animal. Blue squares, daily medians pre-noise; red squares, daily medians post-noise. Error bars show the standard deviation for each day. **B**: Circles: daily median stable period durations (squares in A) averaged across pre- (blue) and post-exposure (red) days. Error bars, standard deviation. **C**: The change in the stable period duration, defined as average daily median duration of post-exposure days (red circle in **B**) minus average daily median duration of pre-exposure days (blue circle in **B**), is shown for the arsanilate- (n = 3) and noise-exposed animals (n = 17). Filled circles denote a significant change (p<0.05 by t-test) in individual animals. **D**, **E**, and **F**: The same as **A**, **B**, and **C**, but for the proportion of stable time. **G**: Change in the stable period duration against the % change in P2N2 amplitude at 1 week after noise exposure for stimulus level 1 through 5 (from left to right). **H**: Change in the stable period duration against the number of CR+ calyces in the saccule (left panel), utricle (middle), and their average (right). **I**: The same as **H**, but for change in the proportion of stable time.

The change in stable period duration was significantly correlated with % change in the VsEP P2N2 amplitude (stim level 5: r = −0.63, p = 0.021; stim level 4: r = −0.66, p = 0.015; stim level 3: r = −0.79, p = 0.0013; stim level 2: r = −0.68, p = 0.011; Figure 4G). The change in stable period duration was also correlated with the number of CR+ calyces in the utricle with marginal significance (r = −0.45, p = 0.065; Figure 4H). Similarly, the change in the proportion of stable time after noise was significantly correlated with the average number of CR+ calyces in the utricle and saccule (r = −0.51, p = 0.036; Figure 4I). Thus, animals that had larger noise-induced dysfunction, as assessed by VsEP amplitude reduction and reduced CR+ calyces in the otoliths, also showed larger increases in the duration of head motionlessness after noise exposure.

In contrast to noise exposure, all 3 animals exposed to arsanilate showed decreased duration of head stability/motionlessness after lesion. There were significant decreases in the proportion of stable time for all 3 animals (Figure 4F), which was an expected result for animals with peripheral vestibular lesions.

### The speed of the animal’s rotational head motion increased or decreased depending on the degree of noise-induced dysfunction

Next, we examined the speed of the rats’ voluntary head motion. We computed the root-mean-square (RMS) of the angular velocity measurements from the x- (roll), y- (pitch), and z- (yaw) axis gyros of the head sensor while the animal’s head was in motion for any duration longer than 0.5 second. Because noise and arsanilate lesions affect active period duration (and stable period duration, see Figure 4), we transformed the RMS values into z-scores using the pre-exposure mean and standard deviation of RMS speeds calculated within small bins of duration. In this way, we minimized the confounding effects of active period duration on RMS angular velocity.

Figure 5A shows the z-score-transformed RMS pitch-axis angular velocity over experimental days in a representative animal exposed to noise. Angular velocity of animals’ head motion was inherently variable; we observed a large variability between segments of motion within a day (error bars) and large day-to-day variability in the daily median RMS (blue squares for pre- and red squares for post-exposure). Nonetheless, most of the daily median RMS values after exposure were below the pre-exposure average (black dashed line), so there was a significant reduction in the daily median RMS pitch-axis angular velocity following noise exposure (p = 0.0055 with Wilcoxon’s rank-sum test, Figure 5B).

**Figure 5:**
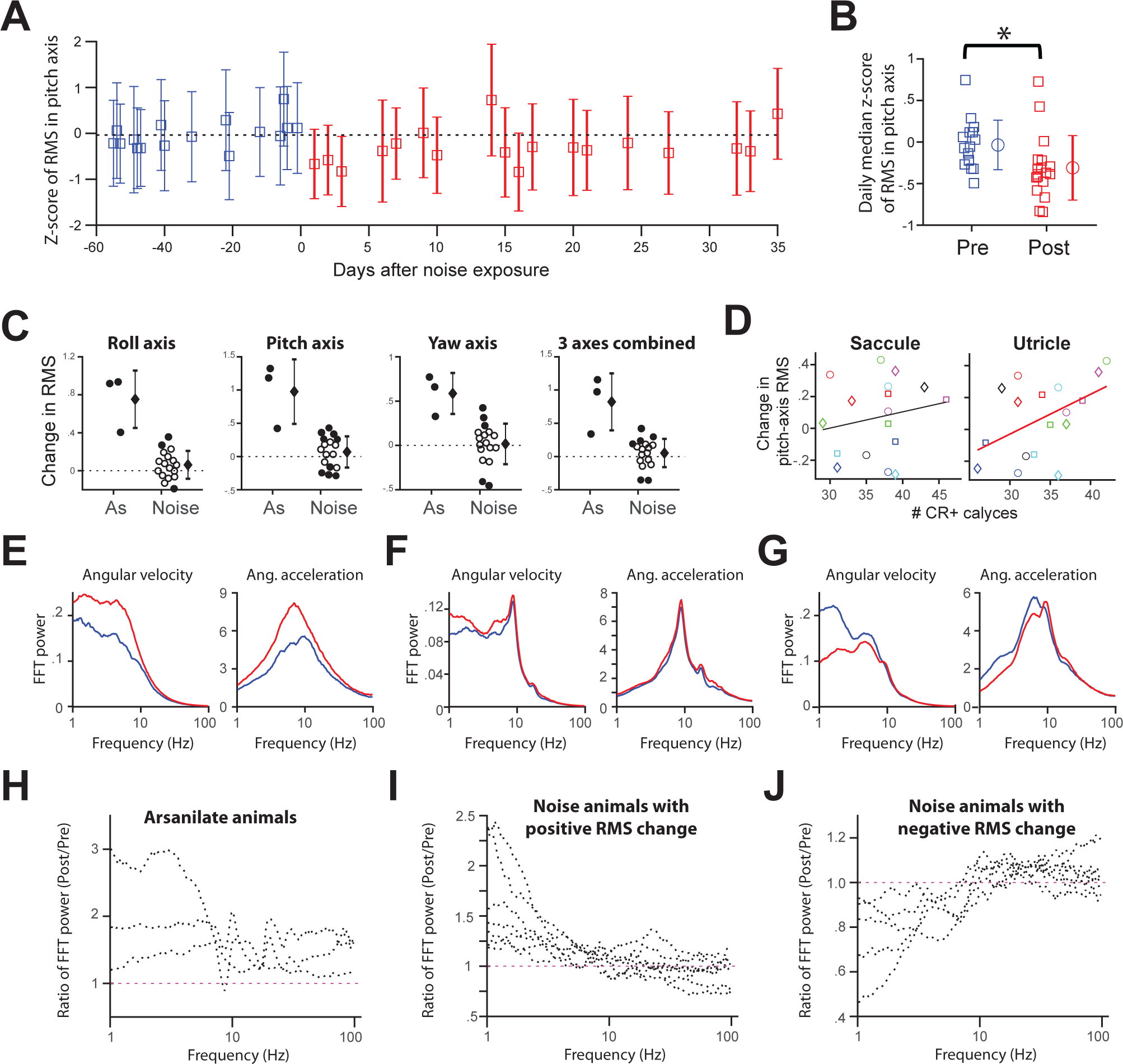
Noise exposure altered the speed of head rotational motion in pitch in some animals. **A**: Speed of pitch-axis rotational motion plotted against days before or after noise exposure. RMS speeds were calculated from pitch-axis angular velocity during any segment of motion longer than 0.5 sec, then transformed into z-scores based on mean and standard deviation of RMS speeds in narrow bins of active period durations from pre-exposure dates. As in Fig. 4, blue symbols denote pre-noise and & red post noise exposure medians. **B**: daily median zscore of RMS speeds (squares) averaged across pre- and post-exposure days. The asterisk denotes significant difference (p < 0.05, t-test). **C**: Changes in z-score transformed RMS angular velocity in roll-axis, pitch-axis, yaw-axis, and vector-average of all 3 axes (from left to right panels) are shown for arsanilate- (n = 3) and noise-exposed animals (n = 17). Change is defined as average daily median z-score transformed RMS of post-exposure days (red circle in **B**) minus that of pre-exposure days (blue circle in **B**). Filled circles indicate statistical significance for individual animals. **D**: Correlation between the change in RMS of the pitch-axis rotational motion and number of CR+ calyces in saccule (left panel) and utricle (right). **E**: Frequency profiles of y-axis rotational motion of before and after arsanilate exposure was obtained by Fourier transform of angular velocity (left panels) and angular acceleration (right panels) when the rat’s head was in motion for durations > 0.5 sec. The median power spectrum before (blue) and after (red) exposure is shown for a representative animal exposed to arsanilate. **F**: the same as **E**, but for a rat whose pitch-axis RMS velocity increased following noise exposure. **G**: the same as **E**, for a rat whose pitch-axis RMS velocity decreased after noise exposure. **H**: The ratio of the post- to the pre-noise power spectrums for 3 animals exposed to arsanilate. **I** and **J**: The same as **H**, but for noise exposed animals with increases (**I**) or decreases (**J**) in RMS y-axis velocity (p < 0.10) following noise.

Out of 17 animals exposed to noise, we found 4 animals with a significant decrease in the daily median RMS pitch-axis angular velocity, and 5 animals with a significant increase (Figure 5C). For the roll- and yaw-axis motion, we found smaller numbers of animals with significant increases or decreases (Figure 5C). When the RMS values from all 3 axes were combined by vector averaging, 4 animals showed a significant increase in overall head speed after noise, while 2 animals showed a significant decrease (right most panel on Figure 5C). Because of these opposed changes following noise exposure, the population-average was diminished to ∼0 for all directions. This contrasts with the changes following arsanilate exposure, where all 3 animals showed a significant increase in head angular velocity in all 3 directions (Figure 5C).

We found a significant correlation between the change in the RMS pitch-axis angular velocity and the number of CR+ calyces in the utricle (r = 0.49, p = 0.048, Figure 5D), but not in the saccule. Some animals with a greater loss of CR+ calyces in the utricle decreased pitch-axis motion speed, while animals with a lessor loss of CR+ calyces increased it. The arsanilate-exposed animals also showed an increase, although the effect size is much larger. Our interpretation of this result is that noise exposure degrades head stability. Together with the results shown in Figure 3, we hypothesize that animals with larger noise-induced damage employ adaptive strategies to reduce both the time duration and the speed of head motion to mitigate head instability brought on by noise exposure.

Examination of the frequency profile of the animal’s head motion before vs after noise exposure hints at a possible stiffening of the animal’s neck musculature as an adaptive strategy to decrease the speed of the y-axis rotational motion. The power spectrum of the pitch-axis angular velocity (left panel, Figure 5E) shows a shallower slope below 5 Hz and a steeper slope in 10 ∼ 30 Hz range. The shape of the power spectrum generally agrees with a previous study, which has reported the power spectrum of rotational head motion deviates from a power law likely due to the biomechanical filtering (Carriot et al., 2014, 2017; Zobeiri et al., 2021). Indeed, when angular velocity is converted into angular acceleration by differentiation, the power spectrum for angular acceleration of the pitch-axis head motion shows a unimodal distribution with a peak just below 10 Hz for all animals tested (see right panels in Figure 5E∼G). In animals who showed head bobbing, the frequency of bobbing coincides at that frequency (Figure 5F). These data suggest that ∼10 Hz represents a resonance frequency for rotational motion of a rat’s head.

In noise-exposed animals with a significant decrease in RMS pitch-axis angular velocity, the power at frequencies below 10 Hz decreased for all animals (Figure 5J). Intriguingly, the power at frequencies above 10 Hz increased in all but one animal. One interpretation of this result is that these animals have stiffened neck musculature, effectively reducing the overall speed of head motion, yet slightly raising the power at higher frequencies (> 10 Hz) following noise exposure.

### Noise-exposure alters animals’ resting head orientation

Lastly, we present how the animal’s head orientation, roll and pitch (Figure 6A and 6H, respectively), changed following noise or arsanilate exposure. Head orientation was calculated from the linear accelerometer measurements whenever the animal’s head was motionless for > 0.1 sec (see Methods for the derivation of roll and pitch angles). Figure 6B shows a scatter plot of a noise-exposed animal’s roll orientation against stable period duration (blue dots for pre- and red dots for post-exposure). Figure 6C shows the mean and standard deviation of roll orientation for pre- (blue line) and post-exposure (red line) as a function of stable period duration. The center of the distribution has shifted by ∼5 degree (tilted toward the animal’s left side) and the distribution broadened after noise, regardless of the stable period duration. In the rest of analyses, we only included head orientation from stable periods > 1 sec to examine the ‘resting’ head orientation.

**Figure 6:**
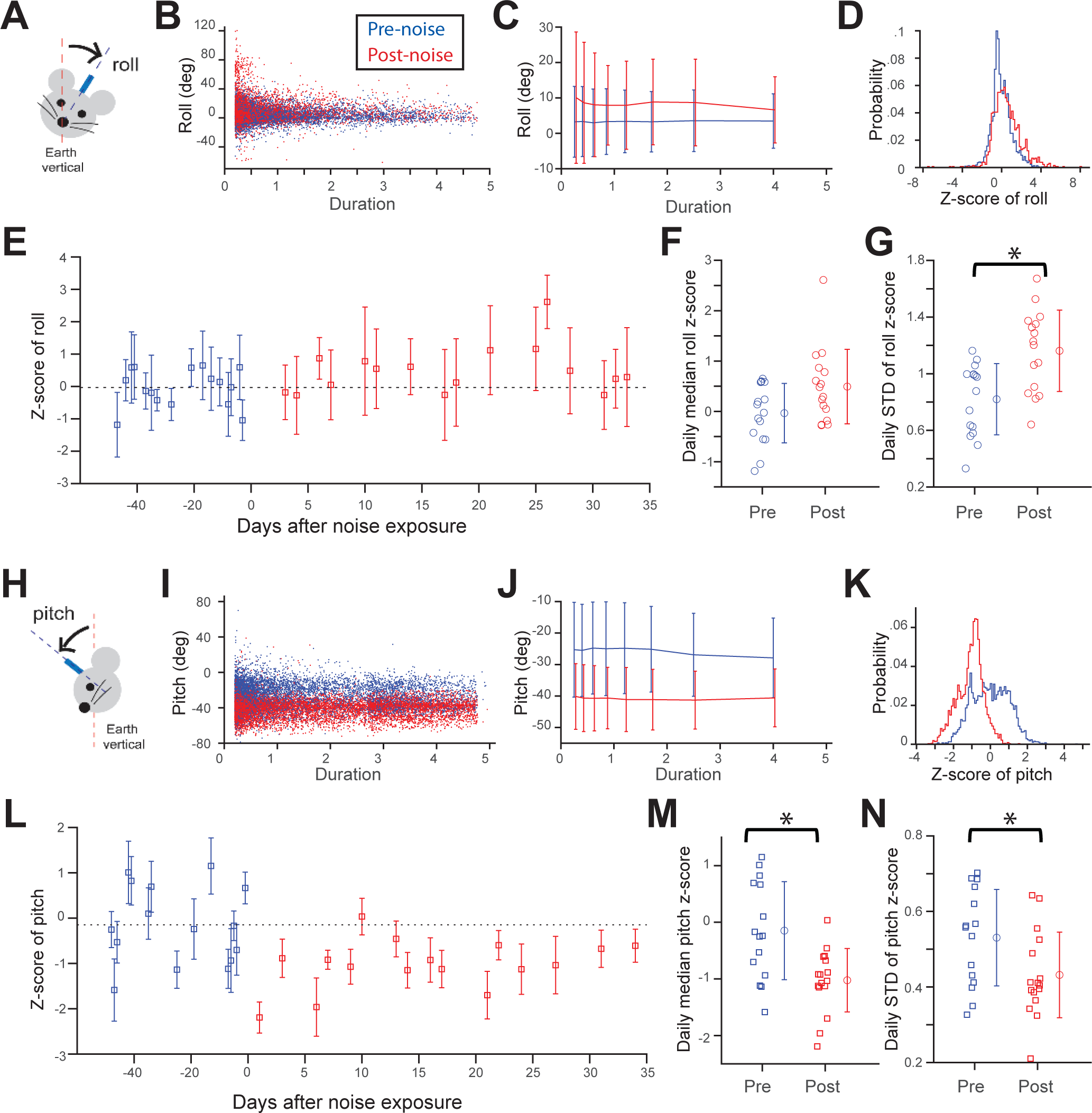
Noise exposure caused changes in head orientation about the roll and pitch axes. A: Roll orientation with respect to gravity and the rat’s head post. B: Distribution of roll angles as a function of motionless period duration for pre- (blue) and post-exposure (red) in a representative rat. C: The mean and standard deviation of roll angle as a function of the duration of motionless period. D: Probability histogram of Z-score transformed roll angles collapsed across durations > 1 sec. E: Z-score transformed roll orientation from motionless periods > 1 sec plotted against days before or after noise exposure. Squares: daily median z-scores of roll; error bars: daily standard deviation. F: Daily median z-score of roll (squares) averaged across pre- and post-exposure days: blue circle, pre-, red circle, post-exposure, error bars, standard deviation. G: The daily standard deviation of z-score transformed roll (squares) averaged across pre- and post-exposure days. Asterisk denotes significance of the mean (p < 0.05 with t-test). H∼N: Same A∼G except these show pitch instead of roll.

The histograms of z-score-transformed roll orientation collapsed across durations >1 sec (Figure 6D) reiterate the shift and broadening in roll distribution after noise exposure. There was a marginally significant increase in the daily median roll orientation (p = 0.08, Figure 6F) and a significant increase in variability within a day (p = 0.0031 with rank-sum test, Figure 6G; see larger red error bars vs. blue in Figure 6E) after noise exposure.

The animal’s pitch orientation was also altered after noise. The plots of pitch orientation as a function of duration on Figure 6I and 6J clearly show a shift and tightening of the distribution regardless of duration after noise exposure in a representative animal. The variability in pitch orientation within each day was significantly reduced after exposure (p = 0.028 with rank-sum test, Figure 6N). The day-to-day variability in median pitch orientation was also reduced, though not significantly (p = 0.105 with variance test, Figure 6M). Both these changes contributed to tightening the overall pitch distribution after noise-exposure in Figure 6K (red histogram). The daily median pitch orientation was also significantly decreased (head lowered toward chest; p = 0.0086 with rank-sum test, Figure 6M).

Of the 17 animals exposed to noise, 4 animals showed a significant change in median roll orientation, with 3 animals tilted to their right side and 1 to the left (Figure 7A). For pitch, 5 animals showed a significant change in median orientation, with 4 animals going downward and 1 upward, following noise exposure (Figure 7D). As a population, the shift in pitch orientation is biased downward, with a statistically marginal significance (p = 0.093 with signed-rank test. Figure 7D), possibly due to gravity. There was a significant correlation between the sizes of the change in median roll and pitch (Figure 7G); an animal with a larger change in median roll orientation tends also to have a larger change in median pitch orientation. Strikingly, the size of change in the median pitch orientation was significantly correlated to the number of CR+ calyces in the saccule (r = −0.67, p = 0.0033 with Pearson’s correlation test; left panel in Figure 7H), but not to the utricle (r = −0.18, p = 0.49; right panel in Figure 7H).

**Figure 7:**
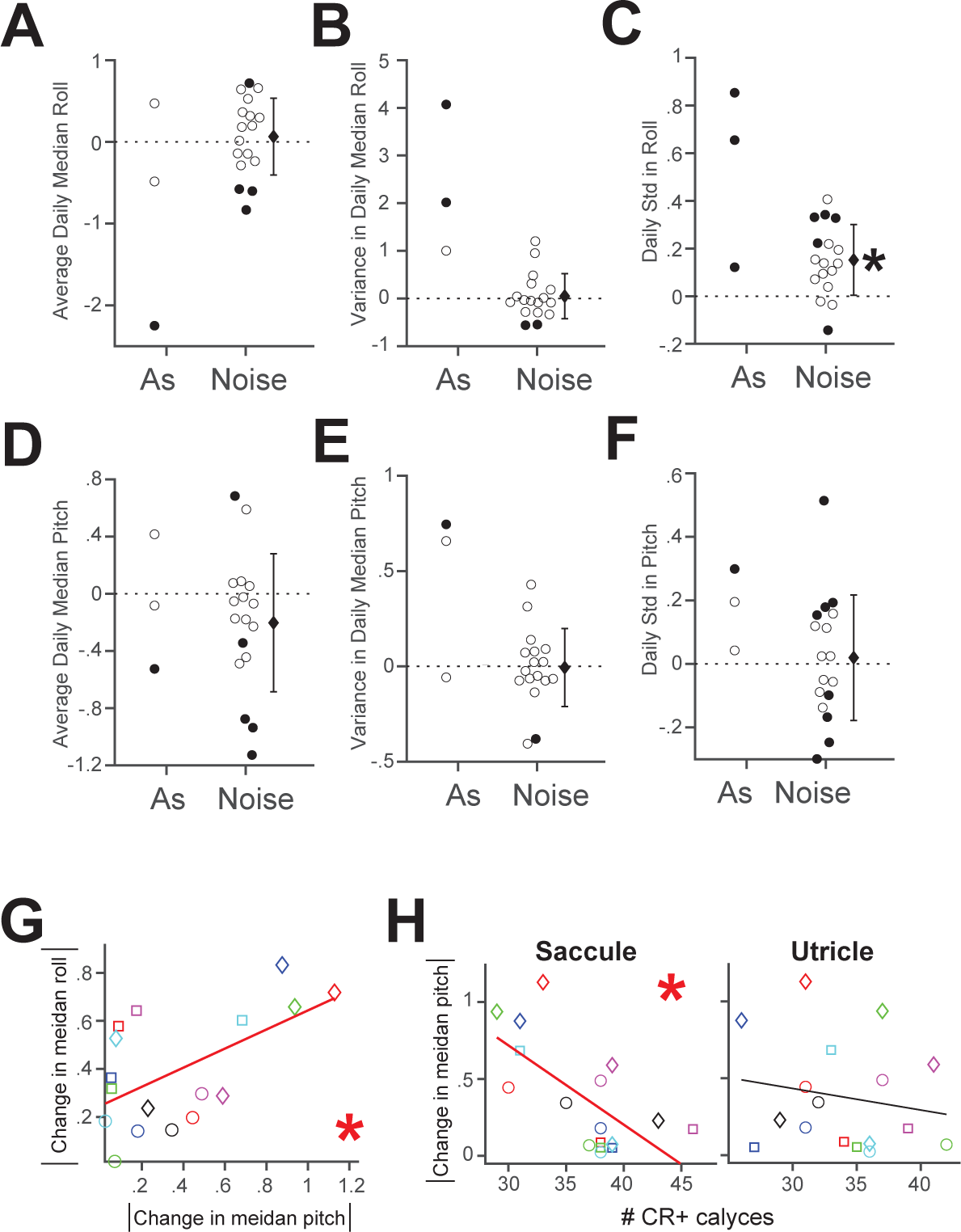
**A** and **D**: Changes in the daily median z-score transformed roll (**A**) and pitch orientations (**D**) for arsanilate (n = 3) and noise-exposed rats (n = 17). Change is defined as the average daily median z-score transformed orientation of post-exposure days (e.g., the red circle in Figure 6F) minus that of pre-exposure days (e.g., the blue circle in Figure 6F). **B** and **E**: Changes in variance of daily median z-score transformed roll (**B**) and pitch (**E**). **C** and **F**: Changes in daily standard deviation of z-score transformed roll (**C**) and pitch (**F**). In **A**∼**F**, filled circles indicate statistical significance in individual animals. **G**: Relationship between absolute values of the change in median pitch and roll. **H**: Relationship between absolute value of the change in median pitch and number of CR+ calyces in saccule (left panel) and utricle (right panel).

## Discussion

### Noise exposure and head stability

Animals with greatest loss of CR+ calyces showed a significant decrease in the speed of y-axis head motion, while those with a lessor loss of CR+ calyces showed significant increases in motion. In addition, animals with larger noise-induced changes in the VsEP and/or the number of CR+ calyces held their heads motionless for longer time periods following noise exposure. Together, we interpret this result as (1) noise exposure was inherently challenging to an animal’s head stability, and thereby manifests as an increase in average head speed in mildly to moderately affected animals, but (2) when the damage was large enough, animals exhibited reduced duration and speed of head motion as a behavioral adaptation to modified vestibular afference.

In patients with vestibular disturbances, a reduction in natural head movements is a common behavioral adaptation because the movement aggravates their vestibular symptoms (Colledge et al., 1994; Heidenreich et al., 2010; Paul et al., 2017). This supports our finding of prolonged head motionlessness in animals with a larger noise-induced damage. Also, an adaptive strategy that includes co-contractions of antagonistic neck muscles to stabilize head upon body perturbations has been reported in elderly subjects (Keshner, 2000). Our result is consistent with the human findings since (1) the animals in our study were also subjected to rapid, horizontal body perturbations that required them to stabilize their unsupported heads and (2) the power of rotational head motion over high frequencies (> 10 Hz) increased following noise exposure in animals which had an overall reduction in RMS speed of motion. Thus, we hypothesize that animals with larger noise-induced damage in the vestibular periphery likely stiffen their neck musculature as an adaptive strategy.

Alternatively, increased power in high-frequency head motion could be interpreted as the functional consequence of the dysfunction of irregular afferents, which are better at encoding natural head movement at higher frequencies than regular afferents (Jamali et al., 2019). Indeed, we found that animals with a delay in the VsEP N1 and P3 latency showed an increased power in high-frequency (> 10Hz), pitch-axis head motion following the noise exposure (Figure 8A, B). The result is in accordance with a study of patients with unilateral vestibular loss, where significant increases in high-frequency head movements were seen when patients performed challenging tasks, such as walking on foam that likely required online updating of an internal model of self-motion (Zobeiri et al., 2021). Because there was no change in the power of low-frequency head movements, for which vision and proprioception could aid stabilization, the increased power of high-frequency head motions in the patients was interpreted to be due to impairment of the vestibular system, which contributes to damping high-frequency head motion.

**Figure 8:**
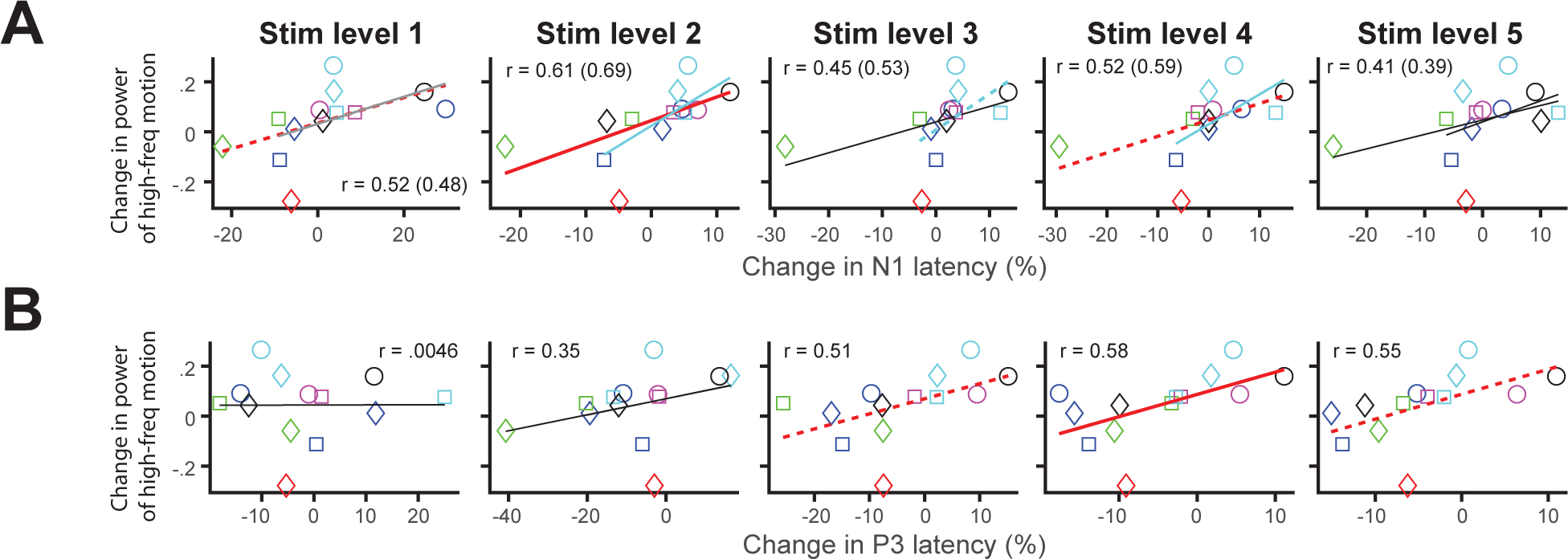
A: The change in the power of high-frequency pitch-axis motion against the change in N1 peak latency at 1 week after noise exposure for stimulus level 1 through 5 (from left to right panel). The power was integrated across frequency > 10 Hz from Fourier transform of pitch-axis angular velocity. B: the same as A, but against the change in P3 peak latency at 4 week after noise exposure. In A, blue lines show the linear fit excluding an outlier data point (denoted with green diamond), and the correlation coefficients with Pearson’s correlation test are shown in prentices.

### Noise exposure alters animal’s resting head orientation

Noise exposure changed the animal’s median head orientation in both roll and pitch. One interpretation is that the change is specific to the behavioral context of the present study, because head orientation was measured during the inter-trial-intervals of imposed horizontal rotations. As mentioned above, an adaptive strategy of co-contraction of antagonistic neck muscles during body perturbations could contribute to a change in head orientation following noise exposure. Resting head orientation during natural movements will need to be examined to confirm this interpretation.

An alternative interpretation is that noise exposure degrades the directional estimate of gravity. Otolith regular afferents likely encode the best estimate of static head position (Jamali et al., 2019); however, afference from the semi-circular canals must also be combined to distinguish head tilt re gravity from linear translational motion (Angelaki et al., 1999). Therefore, the change in resting head orientation following noise exposure suggests that otolith regular afferents and/or canal afference is compromised. Indeed, results from previous studies are consistent with this conjecture (Fetoni et al., 2009); for example, exposure to a broadband white noise at 116 dB SPL for 3 hours significantly reduced the number of the hair bundles on the hair cells in the striolar and extra-striolar regions of the saccules and utricles as well as the anterior and horizontal semicircular canal cristae (Stewart et al., 2016).

We found a significant correlation between the size of the post-noise change in pitch and the relative number of CR+ calyces in the saccule but not the utricle. This points to a strong contribution of saccular irregular afferents to the animal’s resting head orientation regardless of whether it is specific to the behavioral context of the present study. Because the saccules are anatomically more aligned with the earth vertical axis, a greater correlation between saccular damage and the change in an animals’ pitch position is reasonable. In addition, vestibular nucleus neurons that project to the upper cervical cord predominantly receive irregular afferent inputs, whereas those neurons that project to the extraocular muscles (VO and VOC neurons; Cullen et al., 2001) receive higher proportions of regular afferent inputs (Boyle et al., 1992; Highstein et al., 1987). The difference in the ratio of regular vs irregular inputs to neurons involved in vestibulo-ocular vs vestibulo-spinal pathways may reflect the higher inertia of the head as compared to the eyeball, since the large phase-lead exhibited by irregular afferents’ responses to head motion may be necessary to control head position. Thus, irregular afferents, including those that are CR+, are critical to control static and dynamic posture of the head and body, and a change in resting head position may reflect their loss.

### VsEP and non-quantal transmission

We found that animals with a larger reduction in CR+ calyces in the otolith organs showed reduced P1N1 amplitudes and delayed P1 latencies, while those with smaller reductions in CR+ calyces showed slightly enhanced P1N1 amplitudes and earlier P1 peak latencies (Figure 3E, F). In the chicken, the P1 and N1 peaks were suggested to reflect compound action potentials of vestibular irregular afferents based on their resistance to brain stem lesions and a small dependence on body temperature (Nazareth and Jones, 1998). Thus, a reduction in the P1N1 amplitude and delay in the P1 peak latency may represent a delay in irregular afferents’ spike discharges (e.g., first spike latency) and less synchrony in irregular afferents’ discharges.

A very recent study has shown strong evidence that vestibular irregular afferents’ short-latency response to bone conducted vibration is generated through non-quantal transmission at the synapse between vestibular type-1 hair cells and calyx-bearing afferents (Pastras et al., 2023). The basolateral membrane of type-1 hair cells is enclosed by a calyceal membrane at close apposition, and there is a large resting membrane conductance due to low-voltage activated K+ channels and HNC channels on both sides of the synapse (Contini et al., 2020, 2017). The electrical coupling due to this arrangement enables hair-cell depolarization by mechano-transduction currents to trigger action potentials in calyx-bearing afferent fibers with extremely short latencies (nearly instantaneously) and without activation of glutamate receptors (Contini et al., 2020, 2017; Govindaraju et al., 2023).

Assuming the earliest components of the VsEP represent irregular afferents’ compound action potentials driven by non-quantal transmission, then delayed and reduced P1 peak seen in animals most affected by noise exposure may be interpreted as reduced speed and less synchrony in non-quantal spike activation following noise exposure. A computational model by Govindaraju et al., (2023) has shown that the height of the calyceal enclosure on the hair cell affects the speed of non-quantal transmission. Retraction of calretinin-positive calyces has been noted following exposure to vestibulo-toxic gentamicin (Sultemeier and Hoffman, 2017), which can lead to greater action potential discharge latencies according to the model. Whether or not noise exposure also causes calyceal retraction is an interesting hypothesis that must be tested in future experiments.

Interestingly, animals with a smaller reduction in CR+ calyces showed slightly enhanced P1N1 amplitude and faster P1 peak latency. This may be due to a possible compensation by the efferent system on the calyx-bearing afferents. A study has shown a dose-dependent enhancement/suppression of VsEP components by the antagonist and activator of KCNQ channels, which are expressed at the calyces and modulated by efferent activity via muscarinic acetylcholine receptors (Lee et al., 2017).

### Noise exposure affects the expression of calcium-binding protein in the calyx

In our study, the saccule and utricle appeared similarly susceptible to noise exposure, at least regarding the changes manifested in the number of CR+ calyces. Otolith irregular afferents, consisting of calyx-only and striolar dimorphic afferents, can respond to air conducted sound with intensity thresholds 80∼100 dB SPL for the saccule and 100∼120 dB SPL for the utricle over the frequency range of noise used in the present study (Curthoys et al., 2016). Their firing rate steeply increases with intensity above threshold, suggesting that a noise intensity of 120 dB SPL as used in the present study caused excessive firing in otolith irregular afferents, leading to noise-induced damage.

We found a striking correlation between reduced numbers of CR+ calyces in the otoliths and the reduction in the VsEP N2P3 amplitude following noise exposure. This points to a dominant role for CR+ (calyx-only) afferents in generation of the VsEP. At the same time, it raises a question of the degree of contribution by irregular dimorphic afferents in generation of the VsEP as well as their susceptibility to noise exposure. Calyx-only afferents in the utricle, on average, have higher irregularity in their firing pattern and larger phase leads to head motion stimulation compared to irregular dimorphic afferents (Goldberg et al., 1990). This may make calyx-only afferents more sensitive to high-frequency changes in linear acceleration during jerks and be a dominant source of the VsEP response. A future examination of the expression of calbindin in calretinin-negative calyces would address the susceptibility of irregular dimorphic afferents to noise.

Our previous report showed that reduction in CR+ calyces occurred without a significant change in the total number of calyces in the saccule following noise exposure (Stewart et al., 2020). This suggests that the reduction in the number of CR+ calyces is due to down-regulation of calretinin in a subset of calyx-only afferents, and not due to a loss of these afferent terminations.

Noise-induced down-regulation of calretinin would alter the dynamics of the intracellular free Ca^2+^ in calyx-only afferents, which in turn would influence the activity of calcium-dependent K+ channels, HCN channel via cAMP, KCNQ K+ channels via calmodulin, and efferent modulation via G-protein coupled mechanism (Holt et al., 2017, 2015; Lee et al., 2017; Meredith et al., 2011; Seseña et al., 2014; Ventura and Kalluri, 2019). The calix-only afferents contain calbindin, calcium pumps, and many mitochondria in addition to calretinin (Lysakowski et al., 2022; Lysakowski and Goldberg, 1997), indicating that their intracellular free Ca^2+^ must be intricately regulated through mechanisms having different spatiotemporal dynamics. We found a robust correlation between the number of CR+ calyces and a change in head stability and orientation. This points to the relevance of calcium regulation for calyx-only afferent function and ultimately to vestibular behavior. Future studies involving intracellular recording of calyx-bearing afferents would be instrumental in revealing the role of calcium regulation on the functional properties of these afferents in the context of noise-induced damage.

## Methods

### Animals

We used adult Long-Evans rats of both sexes (Charles River Laboratories, MA) with weight ranging between 510 ∼ 860 g in males and between 270 to 430 g in females. They were caged in pairs or trios with free access to food and water. The housing facility had a 12/12-hour light/dark cycle, and all experiments were done between 8am-5pm during the light cycle. Twenty rats were the subjects in the behavioral tests, VsEP recordings, and the immunohistological study. Seventeen rats (8 males and 9 females) received noise exposure while three male rats received sodium arsanilate. Sixteen additional rats served as control subjects in the immunohistological study. Two other rats received noise exposure and served in immunohistological study. All procedures were carried out in accordance with National Institutes of Health guidelines and were approved by the Institutional Animal Care and Use Committee at the University of Michigan.

### Noise exposure

Rats were anesthetized with ketamine (University of Michigan B2 Pharmacy, MI and Covetrus, OH) and xylazine (Covetrus, OH and MWI Animal Health, ID). They received 3-octave band noise (530 ∼ 4240 Hz) bilaterally for 4 hours and were given Atipamezole (Covetrus, OH) as reversal after the completion of the exposure. The waveform of the noise stimulus was created using Adobe Audition version 1.5 software (Adobe, CA). The sound was amplified with an amplifier (DA134, Rolls, UT) and delivered through inserted earphones (ER 3C, Etymotic Research Inc., IL). The sound level was calibrated with a fast Fourier transform spectrum analyzer (SR760; Stanford Research Systems, CA) by setting the peak of the frequency spectrum to 120 dB SPL. The SPL of the spectrum analyzer was calibrated with the sound level calibrator (R8090, Reed Instruments, NC). Bruel & Kjaer type 4136 microphone (Bruel & Kjaer Sound and Vibration Measurement, Denmark) was used in calibration.

### Arsanilate exposure

The stock solution was prepared by dissolving sodium arsanilate in 0.9% sterile saline at 150 mg/mL concentration. An animal was given oral administration of Meloxicam and then anesthetized with isoflurane (MWI Animal Health, ID). Under surgical loupe, 20 µl of the stock solution was injected through the tympanic membrane with a Hamilton syringe. The animal was held on its side for 5 min to allow absorption before repeating the injection in the other ear. Because arsanilate is known as a vestibulo-toxic chemical that destroys the epithelium of the vestibular end organs and degrades vestibular behavior ((Hunt et al., 1987; Ossenkopp et al., 1990; Vignaux et al., 2012), we treated arsanilate-exposed animals as a positive control of bilateral vestibular loss.

### Surgical implantation

The rat was anesthetized with isoflurane (MWI Animal Health, ID) and its head was held in a stereotaxic frame. A midline incision was made to expose the bregma and lambda on the dorsal surface of the skull. The stereotaxic positioning was further adjusted so that the Z-axis coordinates of the bregma and lambda were equal. Three anchor screws were placed into the skull. A custom, small bolt was placed vertically against the bregma and cemented with C&B Metabond (Perkell, NY). The attachment of the head bolt to the skull was strengthened with dental acrylic fusing the bolt and the skull via the anchor screws. The incision was sutured around the implantation. The animal was placed on post-operative recovery for 10∼14 days.

### Vestibular short-latency Evoked Potential (VsEP)

VsEP was recorded as described previously (Stewart et al., 2020). Briefly, a rat was anesthetized with ketamine and xylazine and its head was affixed to a shaker (ET-132, Labworks Inc., CA) via the surgically implanted bolt so that the direction of the stimulus aligned with the animal’s naso-occipital axis. Jerk stimuli with 5 different amplitudes (∼ 0.3 to 6.4 g/ms) were produced by applying linear voltage ramps of different slopes and durations to the shaker. Jerk is the rate of change in linear acceleration. The stimulus duration ranged between 0.65 ∼1.35 ms. Electrodes (stainless needles) were placed subcutaneously at the vertex (noninverting), mastoid (reference), and hip (ground). Stimuli were delivered as successive positive and negative jerks (∼200 each) and averaged together to minimize electromagnetic artifacts from the shaker.

### Immunohistochemistry

The number of calretinin-positive calyces was determined as described previously (Stewart et al., 2020). Briefly, animals were given a lethal dose of sodium pentobarbital and perfused with 4% paraformaldehyde in PBS. Otic capsules were removed, immersed in the fixative for 2∼4 hours, and decalcified in 5% EDTA in PBS for 3∼4 days at room temperature. The utricle and saccule were dissected, then blocked and permeabilized in 0.3% Triton X-100, 5% normal donkey serum in PBS with Ca^2+^ and Mg^2+^ for 2 hours at room temperature. The primary antibodies were mouse anti-calretinin (CR; 1:1000, Millipore MAB1568), chicken anti-neurofilament H (NF; 1:1,000, Millipore AB5539) or rabbit anti-tubulin beta 3 (Tubb3, 1:1000, BioLegend Poly18020), and rabbit anti-myosin 7a (Myo7a; 1:100, Proteus Bioscience, PAB 25–6790) or Phalloidin Alexa Fluor 647(1:100, Millipore Sigma A22287). A 40X z-series with 1-μm steps were collected with a confocal microscope (Stellaris, Leica, Germany) at a region of interest (Stewart et al., 2018, 2020). Images were analyzed for the presence of CR+ calyces with MetaMorph software. CR+ calyces are defined as CR-positive, Myo7a- (or Phalloidin-) negative, and NF- (or Tubb3) positive circles of diameter greater than 5 μm that appeared below the cuticular plate of the sensory epithelium.

### Motion sensor

A miniature motion sensor (3-Space™ LX Embedded, Yost Labs Inc., Ohio) was placed on the animal’s head via the surgically implanted bolt. The motion sensor consisted of 3 accelerometers, 3 gyroscopes, and 3 magnetometers along 3 orthogonal axes. The sensor was oriented so that the x-axis is aligned with the midline of the animal, the y-axis with the interaural line, and the z-axis with the animal’s head bolt. The motion sensor was connected to a personal computer through a USB connection. A custom C++ program along with the sensor manufacturer’s API protocols was run on Visual Studio (Microsoft, WA) to record accelerometer and gyroscope measurements on the PC. The sampling rate was 500 or 1000 Hz. The dynamic range was at +/− 2 G for accelerometers and +/− 500 or 1000 ° /s for gyroscopes.

### Behavioral test

The rat and motion sensor were placed in a holding device that restrained the body with harnesses. The rat supported its own weight on 4 limbs and was able to move its head freely. The rat and restraint device were then placed on a horizontal, servo-controlled table in a dark booth. The table rotated about an earth vertical axis. In each trial of the behavioral test, the rotator generated an abrupt velocity step of 25, 50, 75, or 100 °/s amplitude in clockwise or counterclockwise direction. After the velocity-step rotation, the table paused for ∼0.2 sec and rotated back to the starting position in a reverse direction at 10 °/sec. The table then paused for a random period (3 ∼ 5.5 sec) before the next trial. The data in the present study was taken during these inter-trial-intervals (ITIs).

### Determination of the period of head stability and head orientation

The animal’s head was characterized as motionless when the following term, A(t), was less than an empirically determined threshold;

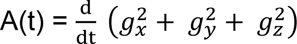, where g_x_, g_y_, and g_z_ are angular velocity in x-, y-, and z-, axis, respectively.

Euler angles of the animal’s head position were calculated for time periods where the animal’s head was motionless, using the following;

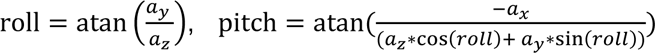, where a_x_, a_y_, and a_z_ are linear acceleration in x-, y-, and z-, axis, respectively.

This pair of Euler angles are based on intrinsic rotations in the ‘yaw-pitch-roll’ order (Kuipers, 2002), which is equivalent to extrinsic rotations in the ‘roll-pitch-yaw’ order.

